# Optimizing CRISPR/Cas9 genome editing in primary human hematopoietic cells to advance studies into HIV biology

**DOI:** 10.64898/2026.08.05.741689

**Authors:** Hong-Ru Chen, Nicole P. Kadzioch, Madeleine Gapp, Hsiu-Hui Yang, Adrian Ruhle, Javier Villamizar Cujar, Thimo Fuchs, Niklas A. Schmacke, Veit Hornung, Roberto F. Speck, Oliver T. Keppler, Manuel Albanese

## Abstract

Defining how human host factors shape HIV-1 infection *in vivo* remains essential for the development of genetically engineered cell therapies. Here, we established a non-viral CRISPR/Cas9 ribonucleoprotein-based platform for efficient single and multiplex gene editing in primary human CD34^+^ hematopoietic stem and progenitor cells (HSPCs). Edited HSPCs retained viability, proliferative capacity, primitive immunophenotypes, and multilineage differentiation potential and supported targeted reporter knock-in independently of the cell source, cord blood (CB), bone marrow (BM), and mobilized from peripheral blood (MPB). Following differentiation, SAMHD1 knockout increased HIV-1 susceptibility of HSPC-derived macrophages, MX2 knockout also increased HIV infection in cells pre-stimulated with IFN-α2a, and CXCR4 knockout blocked X4-tropic HIV-1 infection of HSPC-derived megakaryocytes. We then transplanted CCR5-, SAMHD1-, or non-targeting control-edited HSPCs into immunodeficient mice and challenged reconstituted animals with R5-tropic HIV-1. CCR5 knockout prevented detectable viral spread, validating the model using a clinically relevant dependency factor. In contrast, SAMHD1 knockout accelerated viral dissemination, with earlier plasma viremia and 3.6-fold higher cumulative viremia, although endpoint viral burden was not significantly different from controls. These findings establish transplantation of CRISPR/Cas9-edited human HSPCs as a modular platform for dissecting HIV-1 host-factor function across hematopoietic lineages *ex vivo* and *in vivo* and for evaluating engineered cell-based strategies.

## Introduction

Infection with the human immunodeficiency virus type 1 (HIV-1) remains a major global health challenge despite the success of antiretroviral therapy (ART). Although ART effectively suppresses viral replication, it does not eradicate infection because HIV-1 establishes long-lived latent reservoirs of integrated proviruses that persist in infected cells; consequently, treatment interruption typically leads to rapid viral rebound^1,2^. Activated CD4^+^ T lymphocytes are the main targets for productive HIV-1 infection, whereas resting CD4^+^ T cells are relatively resistant to infection but can harbor latent viruses. Additional cellular reservoirs, including macrophages and dendritic cells, may also contribute to viral persistence in infected individuals^3^. Understanding the cellular and viral determinants that regulate HIV-1 susceptibility in primary human cells is therefore essential for dissecting viral pathogenesis and for identifying host pathways that could be exploited therapeutically.

Human genetic studies have provided strong evidence that host factors can critically shape HIV-1 infection. A landmark example is the naturally occurring 32-base-pair deletion in the gene encoding C-C chemokine receptor type 5 (CCR5), which results in a nonfunctional receptor^2^. The *CCR5-Δ32* mutant helped the identification of CCR5 as the primary co-receptor of HIV-1^4,5^. Its translational relevance was further demonstrated by the long-term remission of an individual HIV-1-infected patient after transplantation with hematopoietic stem cells from a CCR5-Δ32 homozygous donor^6^. These observations illustrate how defining host determinants of HIV-1 susceptibility can inform both basic virology and therapeutic strategy. However, naturally occurring protective mutations are rare and cannot alone support systematic investigation of the many host factors involved in HIV-1 replication, restriction, sensing, and persistence.

On the other hand, SAM domain and HD domain-containing protein 1 (SAMHD1) was identified as a host restriction factor against HIV-1 infection by depleting dNTP pool with its deoxynucleotide triphosphohydrolase activity. Degrading SAMHD1 by Vpx, RNA interference or naturally occurring Aicardi–Goutières syndrome overcame the restriction of reverse transcription and resulted in higher HIV-1 susceptibility in CD4 T cells, dendritic cells, monocytes and macrophages^7–10^. Interferon-induced myxovirus resistance 2 (MX2) is another host restriction factor which contributes to interferon-induced anti-HIV-1 activity. *MX2* gene encoded a member of dynamin-like GTPases family, which is localized in nuclear and cytoplasm. MX2 exerts its antiviral activity against HIV-1 by interfering with viral uncoating^11^ and nuclear import^12–14^. However, studies investigating the restriction function of SAMHD1 and MX2 against HIV-1 infection in *ex vivo* or *in vivo* settings remain limited.

Genome-editing technologies provide a powerful approach to interrogate host gene function directly in primary human cells. Zinc finger nucleases, transcription activator-like effector nucleases (TALENs), and Clustered Regularly Interspaced Short Palindromic Repeats- /associated protein 9 (CRISPR/Cas9) systems introduce targeted double-strand DNA breaks that are repaired by non-homologous end joining (NHEJ) or homology-directed repair (HDR). NHEJ can generate insertions or deletions that disrupt gene function^15^, whereas HDR enables precise knock-in (KI) of defined sequences when a donor template is provided^16^. Compared with earlier nuclease platforms, CRISPR/Cas9 is particularly versatile because target recognition is guided by a programmable RNA sequence rather than by engineering a new DNA-binding protein for each locus^16,17^. Among current techniques, electroporation of Cas9 ribonucleoproteins (RNPs), comprised of synthetic guide RNA (gRNA) and Cas9 protein, has very recently emerged as a method for high efficiency editing in different primary cells, including human CD4^+^ T cells^18,19^. It avoids the need of viral transduction and/or stable genomic integration of CRISPR components, thus reducing toxicity and off-target effects while offering safety advantages for potent clinical applications^20^.

*CCR5* genome editing in primary human CD34^+^ hematopoietic stem and progenitor cells (HSPCs) has been demonstrated using different type of gene editors or virus-based methods in HIV-related studies^21–25^. Very recently, genome editing in these cells using CRISPR-Cas9 has also been performed for studying X-linked chronic granulomatous diseases, sickle cell disease and β-thalassemia^26–29^, with CRISPR-edited HSPC therapy for hemoglobinopathies reaching clinical approval^30^. Together, these studies provided proof-of-principle that individual genes can be genetically manipulated in hematopoietic CD34^+^ precursor cells, followed by successful xenotransplantation and functional *in vivo* analyses.

In this study, we build on prior expertise in Cas9 ribonucleoprotein-mediated editing to develop and optimize KO and KI approaches in primary human CD34^+^ hematopoietic stem and progenitor cells (HSPCs). We apply this platform to genes involved in HIV-1 entry, restriction, and interferon-mediated antiviral activity, including CCR5, CXCR4, CD4, SAMHD1, and MX2. By combining efficient *ex vivo* editing with lineage differentiation into macrophages and megakaryocytes, we establish a system for functional analysis of host factors in HIV-1 infection. These findings support the use of edited CD34^+^ cells not only as a technical basis for improved *in vivo* models of HIV-1 pathogenesis, but also as a biologically informative system to define how individual and combined host factors shape viral infection across human hematopoietic lineages.

## Results

### A highly efficient, low cytotoxic, polyclonal gene knockout method for human CD34^+^ HSPCs

As previously described, our lab has established a method for generating highly efficient, polyclonal, multi-gene KOs in primary human resting CD4^+^ T cells by delivering Cas9-gRNA ribonucleoprotein complexes (RNPs)^19^. Based on this study, we attempted to optimize and extend this approach to CD34^+^ HSPCs. Briefly, we first used a single gRNA targeting CD46 for delivering the RNP complex. We evaluated different amounts of RNP complexes, with 4D- nucleofector programs EO-100, EH-100, DZ-100, and the NEON electroporation system. Using the 4D-nucleofector program EH-100, we achieved 59.3% of CD46 KO in CD34^+^ HSPCs, which was comparable with the efficiency obtained in resting CD4^+^ T cells (68.6%) at both genomic and protein levels (Supplemental Figure 1A). This program outperformed all other tested settings (Supplemental Figure 1A). We further aimed to optimize the protocol by adjusting the cell number and by using a combination of two pre-validated gRNAs targeting CD46 as previously described^19^. Following these conditions, we obtained 98% CD46 KO regardless of the number of cells used (Supplemental Figure 1B).

To support the viability of CD34^+^ HSPCs and preserve their stem-like properties after genome editing, we cultured the cells in an optimized medium containing the molecule UM171^31–33^, β- mercaptoethanol, low-density lipoprotein (LDL), cell growth factors, and cytokines to promote cell proliferation and self-renewal. Under these conditions, we monitored cell viability after nucleofection in non-targeting control (NTC)-treated cells or KO for the HIV-1 restriction factor SAMHD1^8,10,34–37^. On top of that, we also compared CD34^+^ cells from different sources including cord blood (CB), bone marrow (BM), and mobilized from peripheral blood (MPB), to assess whether the origin of the cells affected post-treatment viability. Across all tested conditions, CD34^+^ HSPCs showed 70-80% viability post-nucleofection (Figure 1A). We next asked whether the nucleofection affected cell proliferation. To this end, CD34^+^ cells were labeled with carboxyfluorescein succinimidyl ester (CFSE), with activated and resting CD4^+^ T cells included as proliferating and non-proliferating controls, respectively (Figure 1B). No significant difference was observed among CD34^+^ HSPCs wildtype (WT), NTC, or SAMHD1 KO cord blood-derived CD34^+^ cells (Figure 1C).

**Figure 1.**
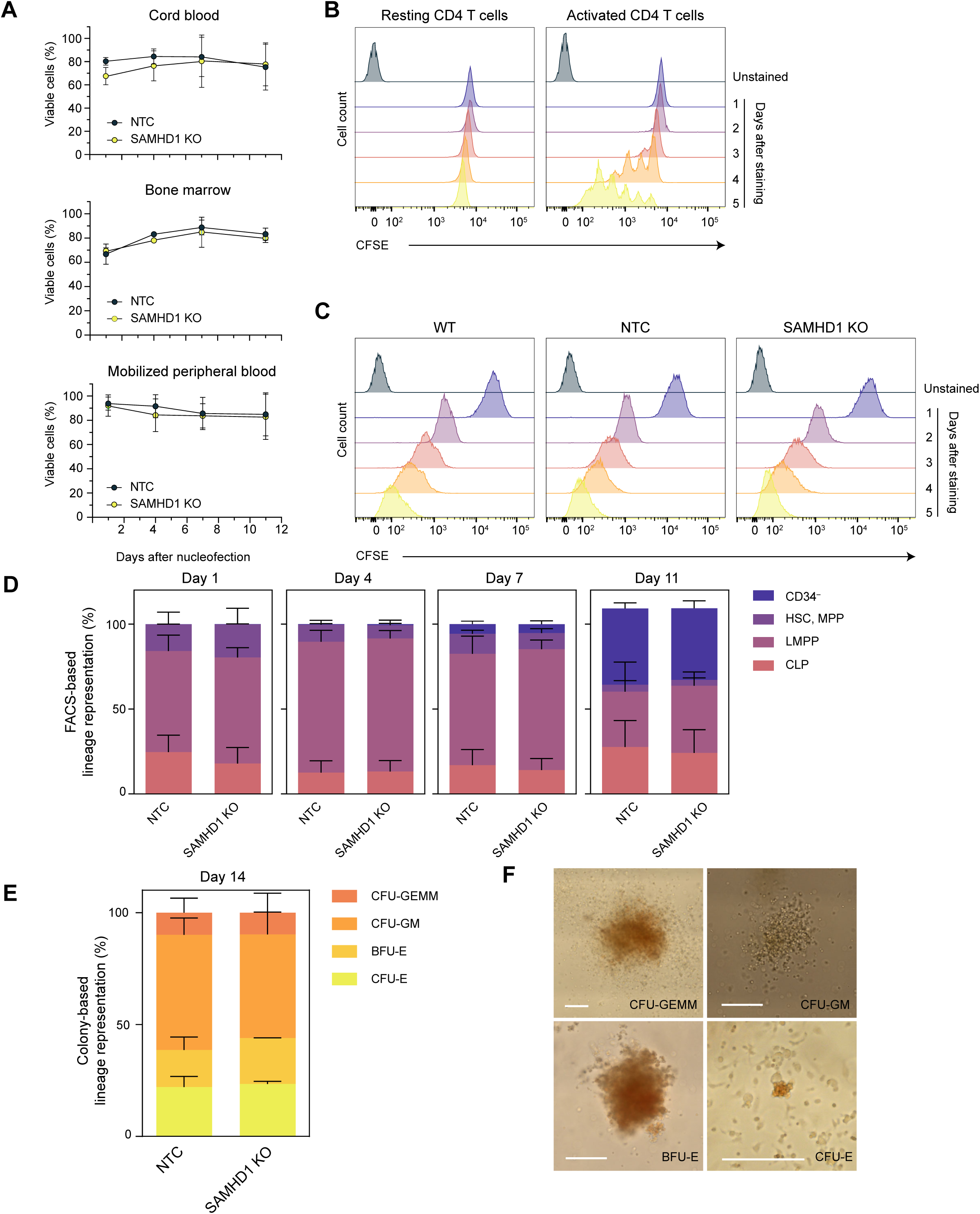
Cell viability, proliferation, and differentiation were not altered by nucleofection with SAMHD1 gRNA in CD34^+^ HSPCs. (A) CD34^+^ HSPCs isolated from cord blood (CB), bone marrow (BM) or mobilized peripheral blood (MPB) were nucleofected with RNPs containing non-targeting control (NTC) or SAMHD1 gRNAs. On day 1, 4, 7,11 post-nucleofection (p.n.), the cells were stained with Live/Dead fixable dead cell stain kit, and the percentage of viable cells was evaluated with flow cytometry. Means ± s.d. are shown (n=2). (B,C) CB CD34^+^ HSPCs were kept untouched or nucleofected with NTC or SAMHD1 gRNAs, followed by staining with CellTrace CFSE one day after nucleofection, and CFSE intensity was acquired with flow cytometry every day until day 5 on CD4 T cells (B), used here as control, or edited) CB CD34^+^ HSPCs (C). One representative experiment out of two is shown. (D) CB CD34^+^ HSPCs were nucleofected with NTC or SAMHD1 gRNAs, followed by staining with CD34, CD38 and CD45RA on day 1, 4, 7, and 11 p.n., and the expression was acquired by flow cytometry. CD34^+^, CD38^-^, CD45RA^-^ cells were classified as HSPC and MPP, CD34^+^, CD38^-^, CD45RA^+^ cells were classified as LMPP, and CD34^+^, CD38^+^ cells were classified as CLP. CD34^-^ cells were differentiated population. Means ± s.e.m. are shown (n=3). (E, F) One day after nucleofection, 1000 CB CD34^+^ HSPCs were seeded in complete MethoCult media in 35 mm dishes and kept in culture for additional 14 days, the images of the full dish were acquired with an Eclipse Ti2 inverted microscope. (E) The colonies were calculated manually. The percentage of each colony with means ± s.e.m is shown (n=3). (F) The representative images of each colony from NTC are shown. Bars represent 200 µm. CFU: colony forming unit; GEMM: granulocyte, erythrocyte, monocyte, megakaryocyte; GM: granulocyte, monocyte; BFU-E: burst forming unit erythroid; CFU-E: colony forming unit erythroid.

Since stemness and differentiation are the most important characteristics of CD34^+^ HSPCs, we assessed these properties by flow cytometry approach and colony-formation unit (CFU) assay. By using CD34, CD38 and CD45RA, we defined CD34+ CD38- CD45RA- cells as hematopoietic stem cells (HSPC) and multipotent progenitor (MPP), CD34+ CD38- CD45RA+ cells as lymphoid-primed multipotent progenitor (LMPP), CD34+ CD38+ cells as committed progenitors and CD34- cells as a differentiated population^38,39^. Over 90% of the initial CD34^+^ cell population maintained its stemness for 7 days in *ex vivo* culture with UM171 medium, with no significant difference between NTC and SAMHD1 KO cells (Figure 1D). CFU assay showed that after nucleofection, the cells were able to differentiate into erythroid colonies BFU-E and CFU-E, granulocyte and monocyte colonies CFU-GM and undifferentiated mixed colonies CFU-GEMM (Figure 1 E and F), which is in line with previous studies^40,41^. No significant difference was observed between NTC and SAMHD1 KO cells, showing that SAMHD1 did not affect the viability, proliferation, and differentiation of the CD34^+^ HSPCs^42^. To validate the SAMHD1 KO efficiency, after nucleofection, CD34^+^ HSPCs (derived from CB, BM, or MPB) were kept in expansion medium containing UM171 for 7 days, and SAMHD1 KO efficiency was evaluated at the genomic level (Illumina Miseq). The result showed a nearly complete loss of SAMHD1 (93.8% to 98.95%) (Figure 2A). Because editing efficiency and post-nucleofection viability were comparable across CB-, BM-, and MPB-derived CD34^+^ HSPCs, these data indicate that our method performs independently of cell source. Therefore, unless otherwise stated, subsequent characterization was performed mainly using CB-derived CD34^+^ HSPCs.

**Figure 2.**
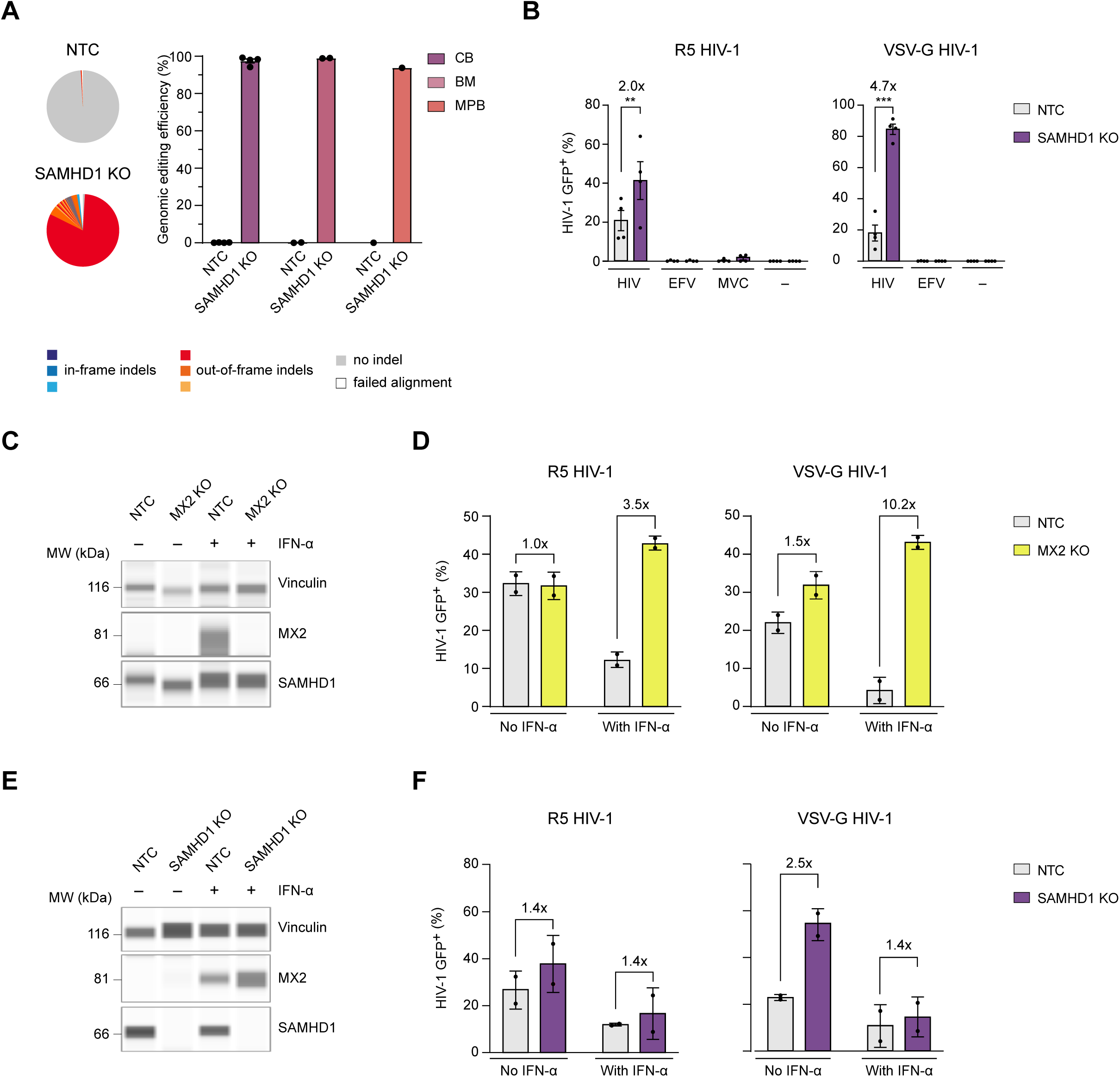
The efficiency of single gene-KO and testing the functionality with HIV-1 infection. CD34^+^ cells were nucleofected with NTC, SAMHD1, or MX2 RNPs, followed by differentiated into monocyte-differentiated macrophages (MDMs). (A) The results of KO efficiency were evaluated by Illumina MiSeq, and the representative pie chart from BM CD34^+^ HSPCs is shown on the left. Means ± s.d. are shown (CB, n=4; BM, n=2; MPB, n=1). (B) Edited CB CD34^+^ HSPCs- differentiated MDMs were infected with R5 HIV-1 or VSV-G HIV-1, with or without Efavirenz (EFV) or Maraviroc (MVC). The percentage of infection was assessed by flow cytometry 3 days post-infection. Means ± s.e.m. are shown (n=4). Asterisks indicate statistical significance by two-way ANOVA; *P* values were corrected for multiple comparison (Šídák). (C, D, E, and F) Edited CD34^+^ cells with NTC or MX2 RNPs (C, D), or with NTC or SAMHD1 RNPs (E, F), were differentiated into MDMs and treated with or without IFN-α2a 10 U/ml for one day. After removing IFN-α2a from culture medium, the cells were subsequently infected with R5 HIV-1 or VSV-G HIV-1. 3 days later, the percentage of infection was assessed by flow cytometry (D, F). The lysate of uninfected cells was collected 3 days after IFN-α2a was removed, and protein level of vinculin, MX2 (C) and SAMHD1 KO (E) were evaluated with JESS capillary Western blotting. Means ± s.d. are shown (n=2). \*\**P* ≤0.01, \*\*\**P* ≤0.001, n.s.: not significant.

### SAMHD1 and IFN-α-Induced MX2 Restrict HIV-1 Infection in CD34^+^ HSPC-Derived Macrophages

After expansion, the resultant CD34^+^ HSPCs were subsequently expanded and differentiated into monocytes in culture with monocyte differentiation medium. One week later, CD14^+^ cells were enriched with CD14 magnetic beads and further differentiated into monocyte-derived macrophages (MDMs). The functional tests were carried out on the MDMs challenged with R5-tropic HIV-1 and Vesicular Stomatitis Virus-Glycoprotein (VSV-G)-pseudotyped HIV-1. As expected, SAMHD1 KO showed higher R5-tropic HIV-1 infection (2-fold), and even higher infection with VSV-G-pseudotyped HIV-1 (4.7-fold) compared to NTC (Figure 2B). In parallel, CCR5 receptor antagonist, Maraviroc (MVC), completely blocked the infection of R5 HIV-1, while the non-nucleoside reverse transcriptase inhibitor, Efavirenz (EFV), blocked the infection of both R5 and VSV-G HIV-1, in both NTC and SAMHD1 KO MDMs (Figure 2B). We further extended this editing approach to another HIV-1 host restriction factor, MX2^43–45^. The expression of MX2 was induced by IFN-α2a in CD34^+^ HSPC-derived MDMs. Targeting MX2 by CRISPR-Cas9 RNPs resulted in a nearly complete KO of MX2 (Figure 2C). In the absence of IFN- alpha pre-stimulation, we did not observe any difference in R5 HIV-1 infection between NTC and MX2-KO cells (Figure 2D), consistent with the lack of MX2 expression under these conditions (Figure 2C). In contrast, IFN-α2a pre-stimulation resulted in a marked increase in viral infection in MX2-KO cells, both after challenge with R5 HIV-1 and VSV-G-pseudotyped HIV-1, with 3.5-fold and 10.2-fold increases, respectively (Figure 2D). In contrast, IFN-α2a pre-stimulation altered neither SAMHD1 expression (Figure 2E) nor the increased susceptibility of SAMHD1 KO MDMs to R5-tropic HIV-1 infection (Figure 2F, left panel). For VSV-G-pseudotyped HIV-1, however, the difference in infection between SAMHD1 KO and NTC MDMs was modestly reduced after IFN-α2 pre-stimulation, from 2.5-fold to 1.4-fold (Figure 2F, right panel). Collectively, these results demonstrate that nucleofected CD34^+^ cells retain their capacity to differentiate into monocyte-derived macrophages in *ex vivo* culture, enabling their use in functional infection assays.

### CXCR4 Mediates X4-Tropic HIV-1 Infection in CD34+ HSPC-Derived Megakaryocytes

HIV-1 genomes have been detected in megakaryocytes isolated from the bone marrow of people living with HIV (PLWH)^46–48^. However, this observation does not establish whether mature megakaryocytes are infected directly or whether viral genomes are acquired by hematopoietic stem and progenitor cells (HSPCs) and retained during their differentiation into megakaryocytes. Only a limited number of in vitro and ex vivo studies have directly examined the susceptibility of megakaryocytes to HIV-1 infection^49–51^. Moreover, genome editing has not yet been applied to megakaryocytes to define the host factors required for viral entry and infection. As a result, we aimed to perform genetic modification on CD34^+^ HSPCs and test the functional consequence in CD34^+^ HSPC-derived megakaryocytes with HIV-1 infection. For ex vivo megakaryocyte differentiation, we adapted a previously described protocol^52^ and cultured the cells for 14 days in megakaryocyte differentiation medium containing recombinant human thrombopoietin (rhTPO), after nucleofection of CD34+ HSPCs followed by a 3-day expansion phase in UM171-containing medium. After 14 days in megakaryocyte differentiation medium, we obtained 89.2% CD41^+^CD61^+^ megakaryocytes as measered by flow cytometry (Supplemental Figure 2A) and confirmed by immunofluorescence, showing mature multinucleated CD42b^+^ CD62P^+^ megakaryocytes (Supplemental Figure 2B). Because HIV-1 infection was performed over 3 days, megakaryocytes were challenged after 11 days of differentiation. At this time, CD41^+^CD61^+^ megakaryocytes represented 63.4% and 70.5% of the NTC and CXCR4 KO cultures, respectively (Figure 3A). CXCR4 editing was performed using two CXCR4-targeting gRNAs, resulting in a near-complete indel frequency as measured by deep sequencing (Figure 3B). This was accompanied by a reduction in CXCR4 surface expression from 52.8% to 0.1%, confirming highly efficient KO (Figure 3C, D). Under these conditions, CD4 expression was not altered in CXCR4 KO megakaryocytes compared with NTC cells, confirming that disruption of CXCR4 did not affect expression of the primary HIV-1 receptor (Figure 3C). CD34^+^ HSPC-derived megakaryocytes were then challenged by spinoculation with X4-tropic HIV-1, R5-tropic HIV-1, or VSV-G-pseudotyped HIV-1. Infection with X4-tropic HIV-1 resulted in 7.3% HIV-1-GFP-positive cells in NTC megakaryocytes, whereas CXCR4 KO blocked infection to a similar extent as treatment with the CXCR4 antagonist AMD3100 (Figure 3F). In contrast, megakaryocytes were not susceptible to R5- tropic HIV-1 infection (Figure 3G). Since the entry of VSV-G HIV-1 is not dependent on the expression of CXCR4, the infection of VSV-G HIV-1 showed similar levels between NTC and CXCR4 KO, while EFV treatment blocked the infection in both cells (Figure 3H). Together, these results demonstrate that CD34^+^ HSPC-derived megakaryocytes support CXCR4-dependent HIV-1 infection and establish genome editing as an effective strategy to define entry-factor requirements in this lineage.

**Figure 3.**
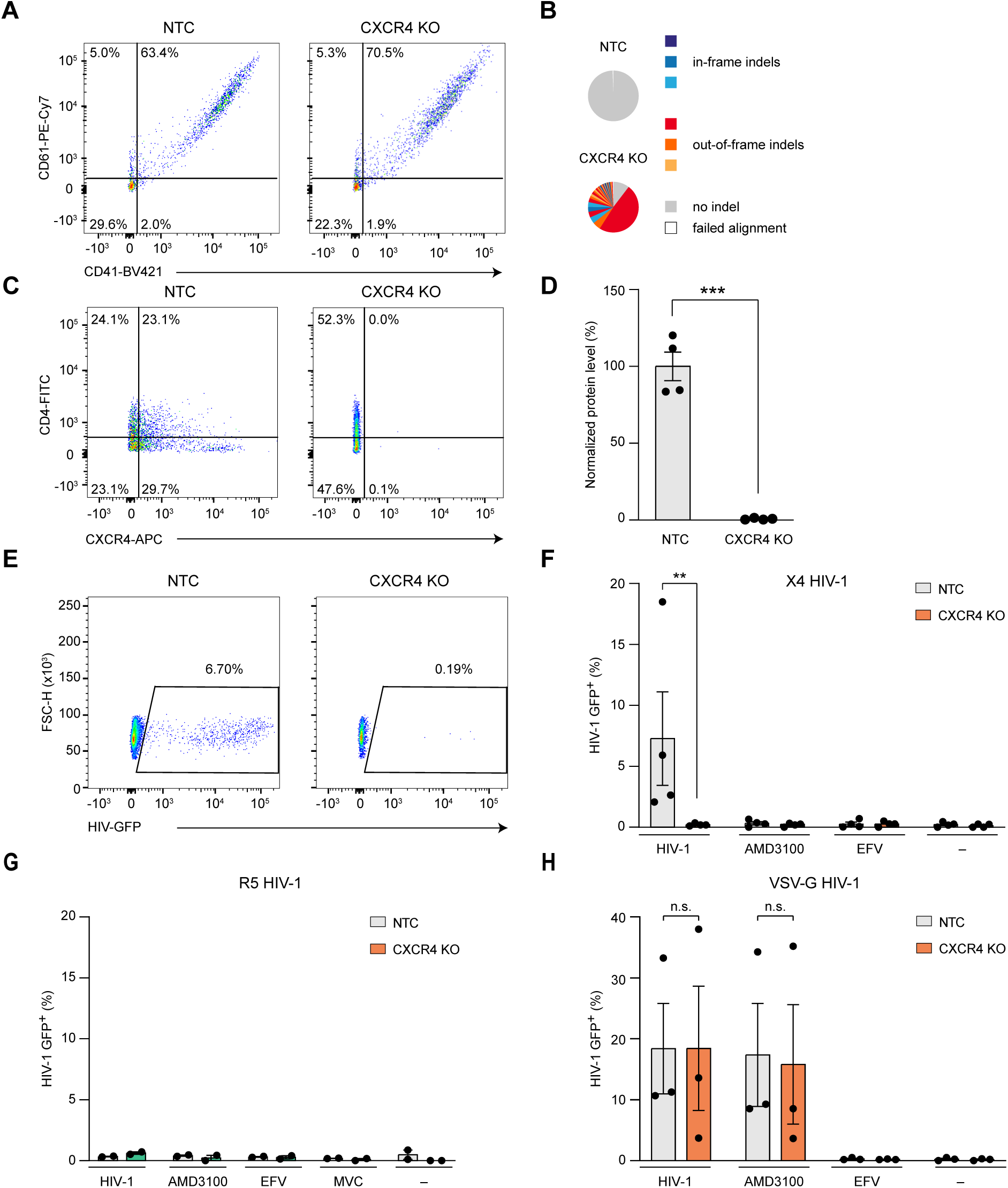
Megakaryocytes differentiated from CD34^+^ HSPCs were susceptible to the infection of X4 HIV-1 via co-receptor CXCR4. CD34^+^ cells were nucleofected with NTC or CXCR4 RNPs followed by differentiating into megakaryocytes in culture. (A) On day 10 post-nucleofection, the expression of megakaryocyte markers CD61 and CD41 was checked with flow cytometry. The CD61^+^CD41^+^ population was defined as megakaryocytes. We proceeded with infection experiments only when CD61^+^CD41^+^ population was over 50%. One representative plot out of four experiments is shown. (B) On day 7 post-nucleofection, cell lysate was collected and sequenced by Illumina Miseq. One pie-chart showing the frequency of indels out of four experiments is shown. (C, D) The expression level of CD4 and CXCR4 on CD61^+^ population was evaluated with flow cytometry 7 days post-nucleofection. One representative plot (C) and the statistics (D) of four experiments are shown. Means ± s.e.m. are shown (n=4). Asterisks indicate statistical significance by two tailed unpaired *t*-test. (E-H) On day 11 post-nucleofection, differentiated megakaryocytes were incubated with X4 HIV-1 (E, F), R5 HIV-1 (G), or VSV-G HIV-1 (H). After stained with megakaryocyte markers CD41 and CD61, the level of HIV-GFP in CD41^+^CD61^+^ population was assessed with flow cytometry. One representative plot (E) and statistics (F) of the infection level of X4 HIV-1 are shown. Means ± s.e.m. are shown (n=4). Asterisks indicate statistical significance by two tailed unpaired *t*-test. The statistics of R5 HIV-1 (G) (n=2) and VSV-G HIV-1 (H) (n=3) infection were also shown with mean ± s.d. (E) or s.e.m (H) and analysed by two-way ANOVA. \*\**P* ≤0.01, \*\*\**P* ≤0.001; n.s.: not significant.

### Multiplex Genome Editing of CD34^+^ HSPCs Enables Functional Analysis of HIV-1 Entry and Restriction Factors

Next, we wanted to extend and apply this protocol for a multi-gene KO approach. We selected 6 genes that are highly expressed in CD34^+^ HSPCs or macrophages (CD4, CD46, PSGL-1, HLA- DR, CXCR4 and SAMHD1). Two gRNAs were selected for each target gene, resulting in 12 different gRNAs in a single RNP nucleofection. Cell viability was assessed 24 hours after nucleofection. Although viability was reduced in the 6-gene KO condition, no significant decrease was observed in the 2- or 4-gene KO conditions compared with NTC cells, and overall viability remained between 67.0% and 79.5% (Figure 4A). On-target frequencies were then evaluated by deep sequencing (Figure 4B). In general, knockout efficiency decreased as the number of targeted genes increased, in contrast to what we previously observed in CD4^+^ T cells^19^ (Figure 4C and D). Consistent with the deep-sequencing data, KO efficiency at the protein level decreased as the number of targets increased, ranging from 51.4% to 97.7% across the tested conditions (Figure 4D). These results show that our genome-editing approach enables multiplex KO in CD34^+^ HSPCs while maintaining cell viability. Because the 4- gene KO condition achieved high KO efficiency for each target, we selected this strategy for subsequent functional characterization in HIV-1 infection assays. We selected 4 critical genes for HIV-1 infection as our KO targets (CD4, CCR5, MX2 and SAMHD1). Deep sequencing revealed that the KO efficiency was higher than 97.6% for the 2-gene KO and 79.7% for the 4- gene KO approach (Figure 5A and B). These results were confirmed by flow cytometry (CD4 and CCR5, Figure 5C) and Western blot analysis (MX2 and SAMHD1, Figure 5D). The lack of CD4 and CCR5 completely protected edited cells from the infection of R5 HIV-1 with or without IFN-α2a pre-stimulation (Figure 5E). In contrast, the infection of VSV-G HIV1 was not dependent on the expression of CD4 or CCR5 as VSV-G HIV enters cells via LDLlow-density lipoprotein receptor^53^. Thus, the infection was not affected between NTC and 2-gene KO regardless whether cells were pre-stimulated with IFN-α2a or not (Figure 5F). In the absence of IFN- α2a pre-stimulation, MX2 protein was not detectable in CD34^+^ HSPC-derived MDMs (Figure 5D). Under these conditions, the 2.5-fold increase in VSV-G-pseudotyped HIV-1 infection observed in 4-gene KO cells primarily reflected the antiviral activity of SAMHD1 (Figure 5F). After IFN- α pre-stimulation, however, the difference in infection between 2-gene KO and 4-gene KO cells increased to 3.2-fold, consistent with the combined antiviral contribution of SAMHD1 and IFN-induced MX2 (Figure 5F). Together, these results demonstrate that our platform supports efficient multiplex gene KO in CD34^+^ HSCs and provides a robust framework for functionally interrogating the individual and combinatorial roles of host-factor during HIV-1 infection.

**Figure 4.**
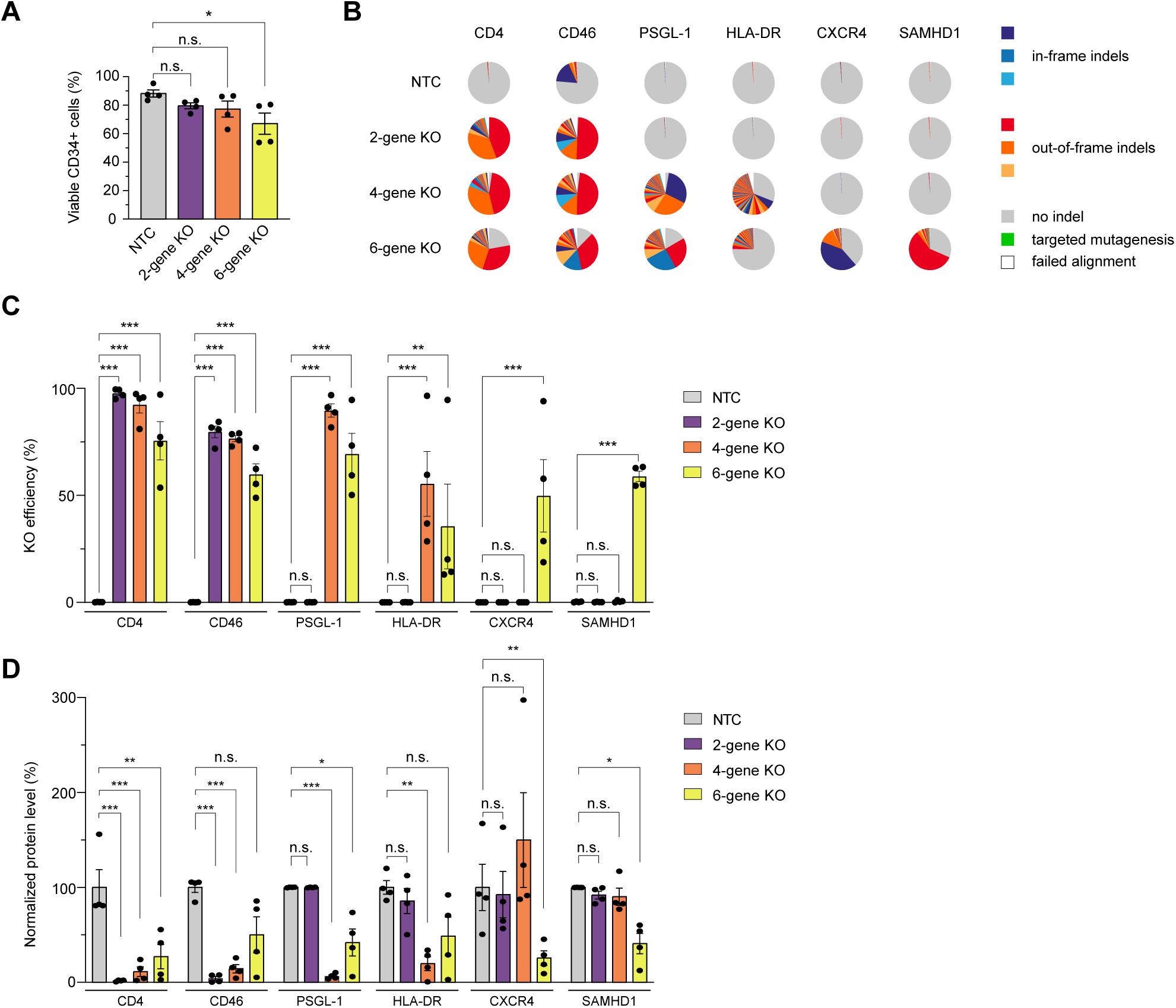
Up to 6 gene-KO could be applied to CD34^+^ HSPCs. CB CD34^+^ cells were nucleofected with NTC RNPs or RNPs targeting simultaneously 2, 4, 6 genes in one single nucleofection. 2- gene KO targeted CD4 and CD46, 4-gene KO targeted CD4, CD46, PSGL-1, and HLA-DR, and 6- gene KO targeted CD4, CD46, PSGL-1, HLA-DR, CXCR4, and SAMHD1. (A) One day post- nucleofection, cell viability was evaluated by flow cytometry. Means ± s.e.m. are shown (n=4). Asterisks indicate statistical significance by one-way ANOVA; *P* values were corrected for multiple comparison (Tukey). (B,C) Seven days after nucleofection, the frequency of indels at on-target sites were assessed with Illumina Miseq sequencing. One representative pie chart (B) and the statistics (C). Means ± s.e.m. are shown (n=4). (D) Ten days post-nucleofection, cells were collected and stained with CD4, CD46, PSGL-1, HLA-DR, and CXCR4, followed by flow cytometry analysis. Means ± s.e.m. are shown (n=4). Asterisks indicate statistical significance by two-way ANOVA; *P* values were corrected for multiple comparison (Dunnett). \**P* ≤0.05, \*\**P* ≤0.01, \*\*\**P* ≤0.001; n.s.: not significant

**Figure 5.**
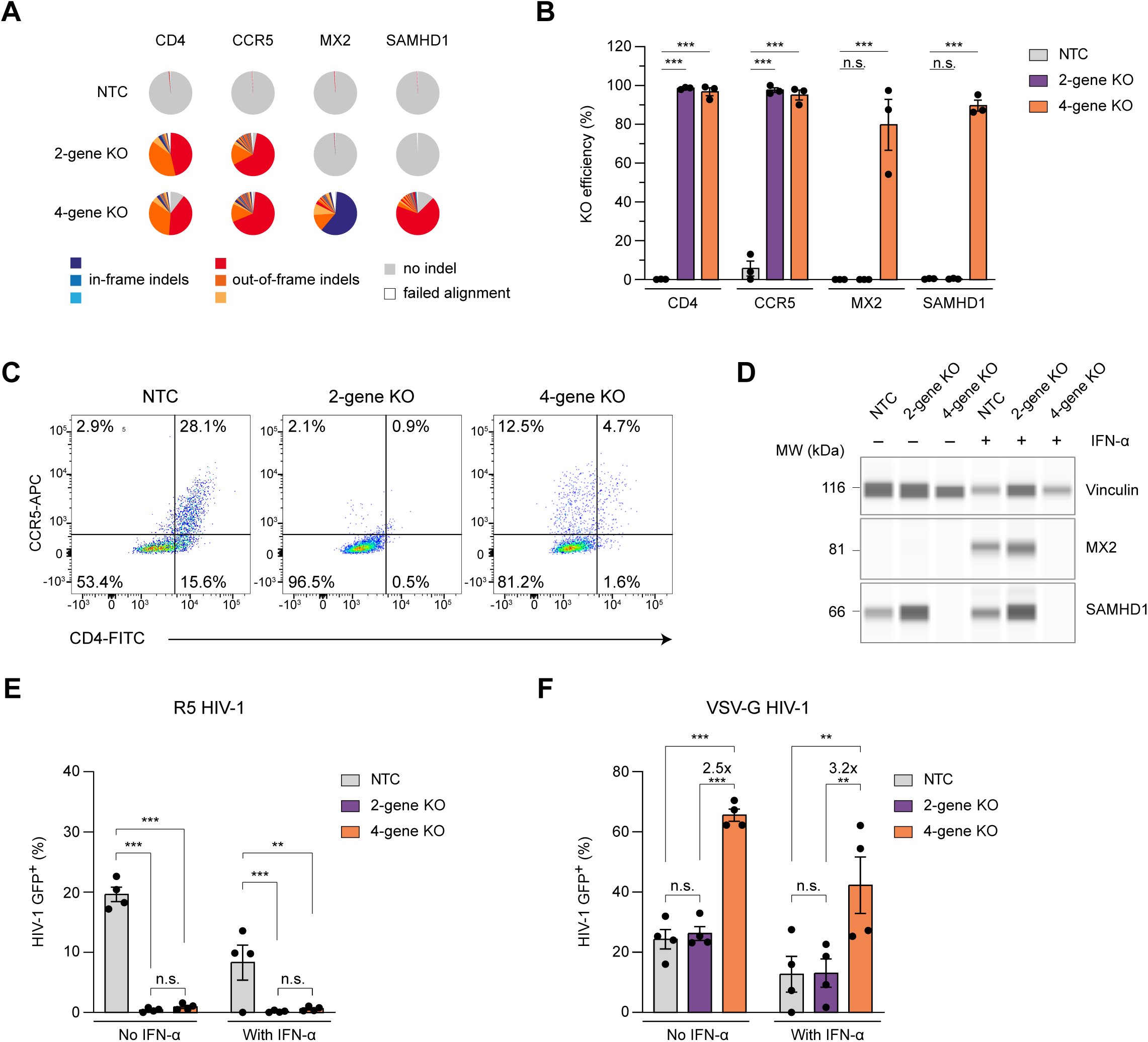
Multiple-gene KO could be applied to functional assays. CB CD34^+^ HSPCs were nucleofected with NTC RNPs or RNPs targeting 2 or 4 genes in one nucleofection. 2-gene KO targets CD4 and CCR5, and 4-gene KO targeting CD4, CCR5, MX2, and SAMHD1. Edited cells were kept in culture and differentiated in macrophages. (A, B) Seven days after nucleofection, cell lysate was collected and sequenced by Illumina Miseq. Pie charts from one representative donor are shown in (A) and statistics (B). Means ± s.e.m. are shown (n=3). Asterisks indicate statistical significance by two-way ANOVA; *P* values were corrected for multiple comparison (Dunnett). (C) Edited cells were kept in expansion medium for 7 days and in monocyte differentiation medium for another 7 days post-nucleofection, followed by enriching CD14^+^ cells with magnetic beads. The expression of CCR5 and CD4 were subsequently checked with flow cytometry prior to further differentiation to MDMs. The plot of one representative donor is shown. (D-F) 21 days post-nucleofection, medium with or without 10 U/ml IFN-α2a was added to differentiated MDMs for one day, and was replaced with complete medium without IFN-α2a on the next day. (D) The lysate of uninfected cells was collected 3 days after IFN-α2a was removed, and protein level of vinculin, MX2, and SAMHD1 were evaluated with JESS capillary Western blotting. (E, F) After removing IFN-α2a from culture medium, the cells were subsequently infected with R5 HIV-1 (E) or VSV-G HIV-1 (F). 3 days later, the percentage of infection was assessed by flow cytometry. Means ± s.e.m. are shown (n=4). Asterisks indicate statistical significance by two-way ANOVA; *P* values were corrected for multiple comparison (Tukey). \*\**P* ≤0.01, \*\*\**P* ≤0.001; n.s.: not significant.

### CRISPR/Cas9-mediated transgene knock-in in CD34^+^ HSPCs

In addition, studying the functionality of a specific gene by disrupting its expression, the CRISPR-Cas9-based genome editing enables the introduction of specific mutation or the insertion of fusion proteins through homology-directed repair (HDR), resulting in precise KI at specific genomic loci. In a previous study, we and other showed that by using Cas9 RNPs and dsDNA, GFP reporter could be targeted at different position in the genome including SAMHD1, Basic Leucine Zipper ATF-Like Transcription Factor (BATF), RAB11A, and CD4 loci in CD4^+^ T cells, resulting in up to 60% GFP-positive cells^18,19^. Since dsDNA showed a certain level of cytotoxicity, we first adjusted the amount of dsDNA for CD34^+^ HSPCs, and found that using 200 ng dsDNA together with Cas9 and a single sgRNA targeting RAB11A resulted in 17.3% GFP positive cells, with >80% viable cells (Figure 6A). After 3 weeks of expansion in UM171- containing medium, the GFP-expressing population could be enriched by flow cytometry-based cell sorting. Control cells, including non-nucleofected cells and cells nucleofected with either sgRNA or dsDNA alone, together with sorted and unsorted GFP-RAB11A-expressing cells, were lysed and analyzed by Western blotting. Control samples expressed only endogenous RAB11A at approximately 25 kDa, whereas a 55-kDa GFP-RAB11A fusion protein was detected with an anti-RAB11A antibody in both sorted and unsorted KI cells (Figure 6B). Using an anti-GFP antibody, the fusion protein was detectable only in the sorted GFP-positive population (Figure 6B). These results show that targeted KI can be efficiently achieved in CD34^+^ HSCs while maintaining high viability, and that KI-positive cells can be further enriched through expansion followed by cell sorting.

**Figure 6.**
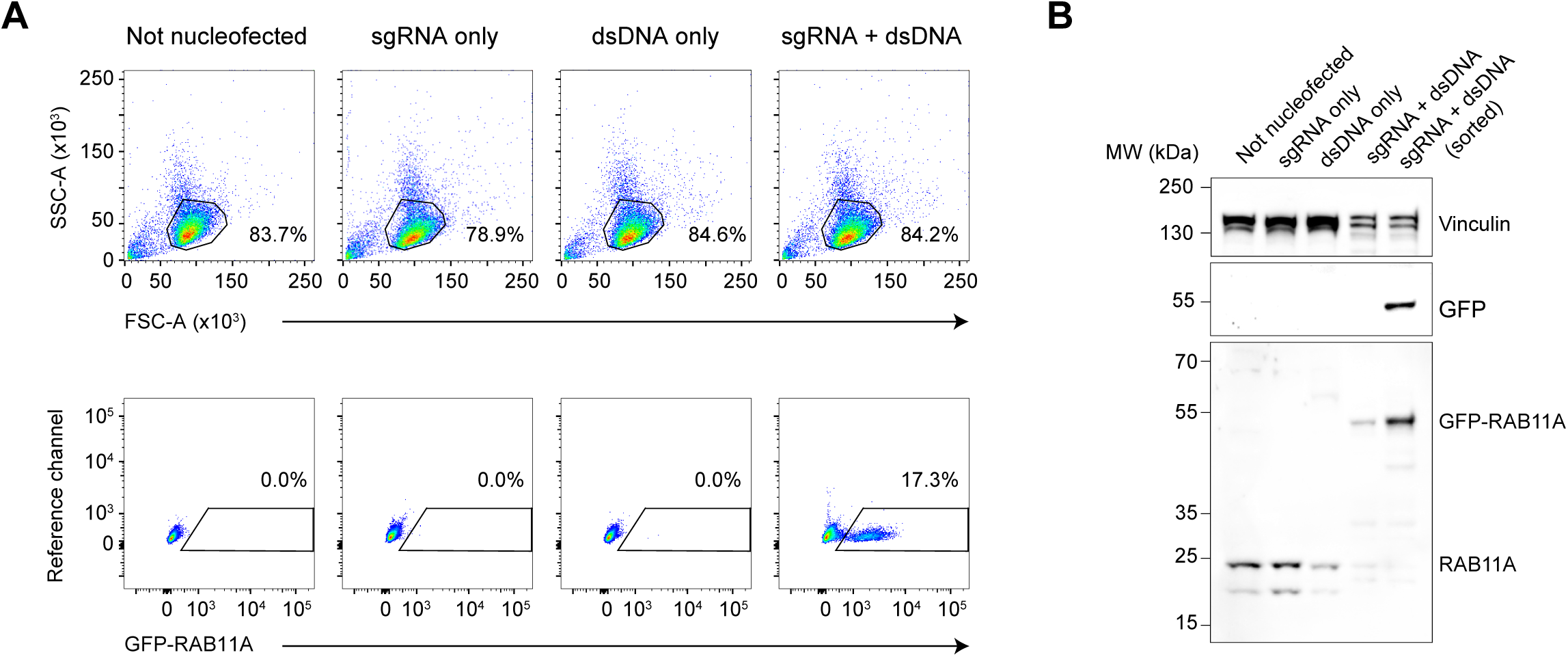
CD34^+^ HSPCs were KI with GFP-RAB11A with high viability and efficiency. CB CD34^+^ cells were nucleofected with RNPs containing sgRNA targeting RAB11A and GFP-RAB11A dsDNA. Cells without nucleofection, nucleofected with only sgRNA, and nucleofected with only dsDNA were used as controls. (A) 7 days after nucleofection, the percentage of GFP-positive cells was assessed by flow cytometry. The living cell population was gated with forward and side scatter and the percentage are indicated (upper panel). Subsequently, the GFP-positive population was gated and the percentage is shown (lower panel). (B) After expanding in culture for 4 weeks, GFP-positive population was enriched by flow cytometry-based cell sorting. Cell lysate from each condition was collected and applied to Western blotting to obtain the protein size and expression of GFP and RAB11A. Vinculin is used as loading control. One donor is shown.

### SAMHD1 Restricts HIV-1 Dissemination in HSPC-Humanized Mice

To determine whether CRISPR/Cas9-edited human CD34^+^ HSPCs could be used not only *for ex vivo* functional assays but also to interrogate HIV-1 biology *in vivo*, we next transplanted edited HSPCs into immunodeficient mice and monitored the resulting humanized animals after HIV-1 challenge (Figure 7A). We selected CCR5 as a benchmark target, because loss of CCR5 should protect from R5-tropic HIV-1 and therefore provides a stringent positive-control system for the model^5,25^. In parallel, we targeted SAMHD1 to test whether the restriction phenotype observed in HSPC-derived macrophages *ex vivo* could be detected in the more complex setting of an infected organism.

**Figure 7.**
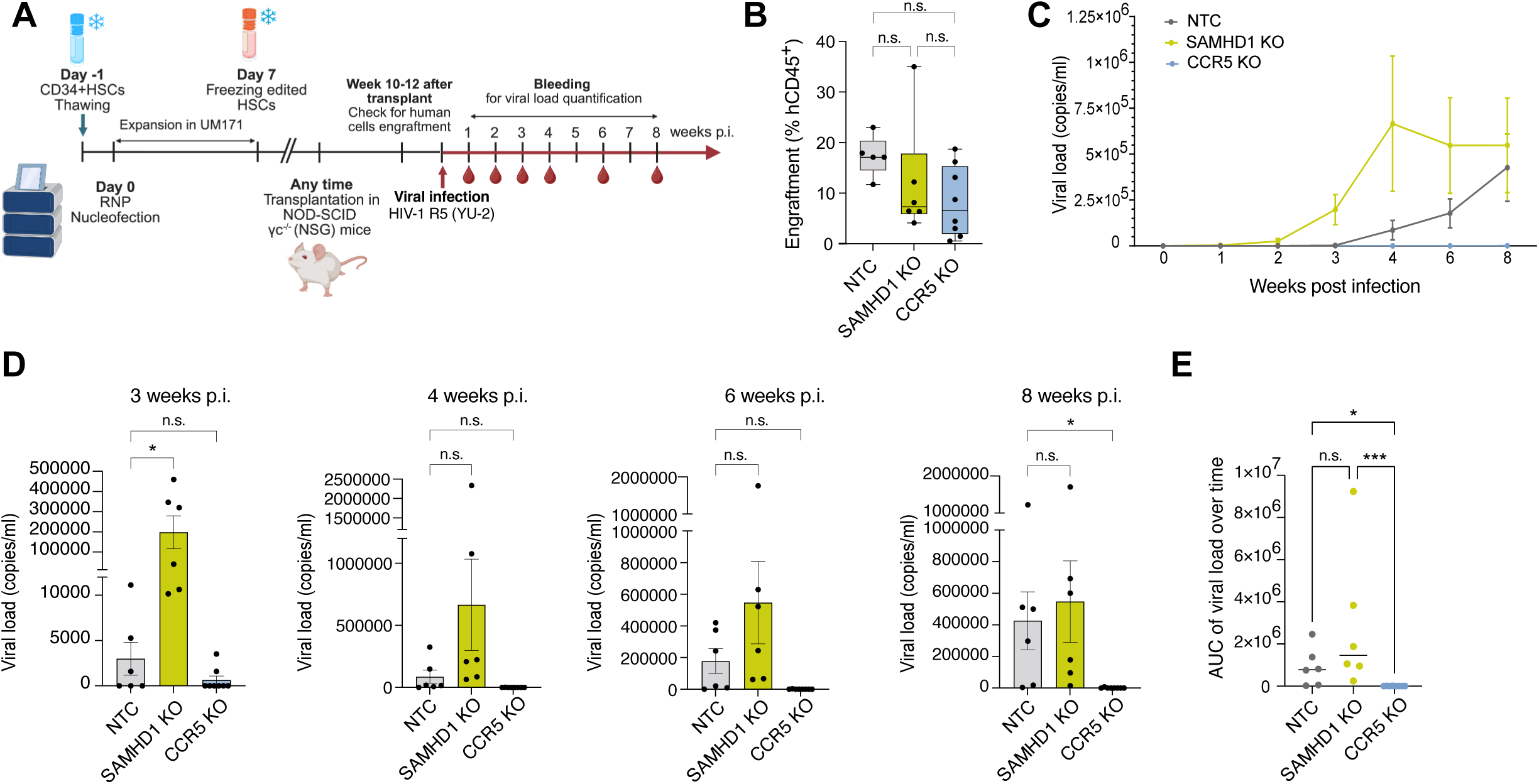
CRISPR-edited CD34^+^ HSPC-derived human immune systems allow functional analysis of HIV-1 host factors *in vivo*. (A) Schematic representation of the experimental workflow. CB CD34^+^ HSPCs were nucleofected with NTC, SAMHD1, or CCR5 RNPs, expanded ex vivo in UM171-containing medium, cryopreserved, and subsequently transplanted into immunodeficient NOD-SCID γc-/- (NSG) mice. After human immune cell reconstitution, mice were challenged with R5-tropic HIV-1 YU-2 and followed longitudinally for plasma viral load. Human cell engraftment was assessed 10-12 weeks after transplantation, followed by HIV-1 challenge and longitudinal blood sampling for viral load quantification. (B) Human cell engraftment was assessed before infection as the percentage of hCD45+ cells in peripheral blood. Individual mice and means ± s.e.m. are shown. (C) Plasma viral load was measured over time after HIV-1 infection in mice reconstituted with NTC, SAMHD1 KO, or CCR5 KO-derived human edited cells. Means ± s.e.m. are shown. (D) Plasma viral load at 3, 4, 6, and 8 weeks post-infection (p.i.) is shown for each experimental group. Individual mice and means ± s.e.m. are shown. (E) Area under the curve (AUC) of plasma viral load over the longitudinal infection course. Individual mice and means ± s.e.m. are shown. Asterisks indicate statistical significance by Kuskal Wallis Test; *P ≤0.05, ***P ≤0.001; n.s.: not significant.

Human CD34^+^ HSPCs were edited with Cas9 RNPs targeting CCR5, SAMHD1 locus, or a NTC gRNA and were transplanted into recipient mice. Following human immune-system reconstitution (Figure 7B), editing was assessed in the human graft, and animals were infected with R5-tropic HIV-1 (YU-2) (Figure 7A)^54^. Plasma viremia was monitored longitudinally over the course of 8 weeks of infection (Figure 7 A).

After R5-tropic HIV-1 challenge, CCR5-KO animals showed no detectable productive viral spread compared with NTC animals (Figure 7C and D). Thus, genetic removal of the principal HIV-1 co-receptor from the HSPC-derived human immune system reproduced the expected protection phenotype *in vivo* and confirmed that the model is sufficiently sensitive to capture host-factor-dependent differences in HIV-1 replication. Following HIV-1 challenge, mice reconstituted with SAMHD1-KO human HSPCs displayed accelerated viral spread relative to NTC control animals. This was evident from an earlier rise in plasma viral RNA at 3 weeks post-infection and a higher cumulative viremia over time, as quantified by the area under the curve (AUC; Figure 7C and E: 480,020 for NTC, 1,714,087 for SAMHD1-KO, and 2,040 for CCR5-KO).

However, viral burden at the endpoint was not significantly different between SAMHD1-KO and control animals (Figure 7D). These data extend the *ex vivo* infection results and demonstrate that SAMHD1 measurably limits HIV-1 propagation within an HSPC-derived human immune system *in vivo*. Importantly, the phenotype was detected in a longitudinal time-course experiment rather than at a single endpoint, supporting the interpretation that SAMHD1 loss accelerates the kinetics of viral dissemination. To our knowledge, this study provided the first direct demonstration in a humanized mouse model that SAMHD1 KO in transplanted human HSCs enhances HIV-1 spread *in vivo*.

## Discussion

In this study, we established a CRISPR/Cas9 RNP-based platform for efficient, polyclonal gene disruption in primary human CD34^+^ HSPCs and showed that edited cells retain key properties required for downstream biological analysis. Across HSPC sources, edited cells remained viable, expanded in UM171-containing culture, preserved primitive immunophenotypes, and retained multilineage differentiation potential. The platform supported single-gene and multiplex knockout, as well as a non-viral KI strategy, and edited HSPCs could be differentiated into macrophages and megakaryocytes for functional HIV-1 infection assays. Most importantly, we extended this approach into humanized mice, where edited HSPCs reconstituted a human immune system that could be challenged with HIV-1 to reveal host-factor-dependent infection phenotypes *in vivo*.

The CCR5 results provide an important benchmark for the model. CCR5 remains the most clinically validated host target in HIV cure research, based on the durable remissions observed after transplantation with *CCR5-Δ32/Δ32* donor cells. Recent preclinical work has sharpened this concept by showing that very high-frequency CCR5 editing in human HSPCs can generate xenografts that are resistant to HIV-1, and that protection declines as the fraction of edited cells falls below high thresholds^25^. Our CCR5-KO humanized mice are consistent with this framework: loss of CCR5 in the HSPC-derived human graft prevented detectable R5-tropic HIV- 1 spread and therefore validates the sensitivity of our system for detecting genetically encoded HIV resistance *in vivo*.

At the same time, the field increasingly recognizes that CCR5 loss alone is unlikely to be sufficient as a broadly applicable HIV cure strategy. R5-tropic virus predominates during transmission, but X4-tropic and dual-tropic variants can emerge, and HIV cure approaches must contend with viral diversity, escape, and reservoir persistence. Recent HSPC-engineering studies have therefore moved toward multilayered resistance strategies, including CCR5 KO combined with KI of fusion inhibitors, restriction factors, or antibody-expression cassettes ^40,55^. Other work has shown that direct disruption of CXCR4 can protect against X4-tropic virus, but may impair hematopoietic engraftment or cell distribution *in vivo*, underscoring the need for alternatives that inhibit X4-tropic HIV without compromising essential CXCR4 biology^56^. In this context, our platform is valuable because it allows host factors and antiviral designs to be tested in primary HSPC-derived lineages both before and after transplantation.

The SAMHD1 humanized-mouse experiment is the central biological advance of our study. SAMHD1 has long been characterized as an HIV-1 restriction factor in myeloid cells and resting T cells^9,10^, and SAMHD1-deficient murine models have been used to examine retroviral restriction and innate immune phenotypes *in vivo*^57^. However, the relevance of SAMHD1 loss in a human hematopoietic system during systemic HIV-1 infection has been difficult to address directly. By generating SAMHD1-KO humanized mice from edited CD34^+^ HSPCs and following HIV-1 replication over time, we show that SAMHD1 restrains viral spread *in vivo* and that this contribution can be underestimated when infection is assessed only at endpoint. However, the absence of a significant difference in viral burden at the endpoint indicates that SAMHD1 loss primarily affected the kinetics of viral spread rather than the final viral burden under the conditions tested. The accelerated kinetics observed in SAMHD1-KO animals indicate that the restriction phenotype is not confined to reductionist *ex vivo* culture systems but remains detectable within the complex reconstituted human immune system.

This result also illustrates the conceptual strength of HSPC editing for HIV biology. Editing a restriction factor, or a putative one, in the stem/progenitor compartment enables the altered genotype to be propagated into multiple hematopoietic lineages, allowing the contribution of that factor to be tested during infection *in vivo*. This differs from transient perturbation in mature cells and from cell-line models, where lineage context, differentiation state, and tissue distribution are incompletely represented. The approach is aligned with recent work showing that RNP-edited human CD34^+^ HSPCs can engraft and maintain edited progeny in humanized mice^58^, and with broader efforts to improve HSPC editing and delivery through all-in-one lentivirus-derived nanoparticles or engineered VLPs^59–61^.

Our *ex vivo* lineage studies broaden the utility of the platform beyond canonical CD4^+^ T-cell infection. HSPC-derived macrophages reproduced expected phenotypes for SAMHD1 and MX2, including increased HIV-1 susceptibility after SAMHD1 knockout and interferon-dependent antiviral activity of MX2. Edited HSPC-derived macrophages remained responsive to type I interferon stimulation, providing a functional platform to study interferon-inducible genes. HSPC-derived megakaryocytes supported analysis of HIV-1 entry dependency, with CXCR4 knockout blocking X4-tropic HIV-1 infection while preserving susceptibility to VSV-G-pseudotyped virus. This is relevant because megakaryocytes and the bone marrow compartment have been implicated in HIV-associated hematological abnormalities, including thrombocytopenia, but remain comparatively underexplored. Our study demonstrates in primary cells that megakaryocytes are susceptible to X4-tropic rather than R5-tropic HIV-1 infection, in agreement with a previous report^62^. The feasibility of CRISPR-edited CD34^+^- derived megakaryocyte systems has also been supported by recent platelet-biology work, which used edited megakaryocytes to interrogate platelet-lineage gene function without compromising megakaryopoiesis^63^.

A practical advantage of this RNP-based approach is that single- and dual-gene knockouts reached near-complete editing across all HSPC sources tested, allowing bulk populations to be used directly for differentiation, infection assays, and xenotransplantation without clonal derivation or selection. Avoiding these steps our protocol limits *ex vivo* culture and cell loss, which can impair HSPC fitness and long-term repopulating capacity^64,65^, while preserving the polyclonal graft composition needed to interpret engraftment and lineage output^66^. Editing remained target- and multiplex-dependent, decreasing to approximately 80% with four targets and further with six, while HDR-mediated knock-in was less efficient than NHEJ-mediated disruption. Enrichment may therefore be beneficial for higher-order multiplexing, refractory loci, or knock-in. Transient AND-gate reporters can enrich correctly edited HSPCs to 80–100% purity and deplete cells with unintended on-target rearrangements while preserving *in vivo* reconstitution^61^. Thus, combining unselected populations for near-complete knockout with enrichment, when editing is limiting, could extend the platform to more complex host-factor combinations.

Humanized mouse models have become valuable and widely used tools for studying HIV infection, antiretroviral therapy responses, and mechanisms of viral persistence and reservoir establishment. However, several limitations should be considered. First, these models do not fully recapitulate the complexity of human immune architecture, lymphoid tissue organization, or long-term ART-suppressed reservoir biology^67^. Second, donor-to-donor variation in HSPC editing, engraftment, and lineage output may influence the magnitude of HIV-1 phenotypes and should be explicitly reported. The low level of human immune cell engraftment achieved in these mice transplanted with edited HSPC may have several possible explanations, including donor-specific characteristics, reduced HSPC fitness after thawing, and potential loss of CD34^+^ HSPCs stemness during *ex vivo* expansion. Importantly, despite the lower overall engraftment levels, the established humanized mouse model allowed the assessment of the functional consequences of HSPC editing *in vivo* within a human immune cell context. Further optimization of HSPC editing strategies and transplantation conditions may improve engraftment efficiency and enhance the application of the model for future studies. Third, our longitudinal design identifies when SAMHD1 loss alters viral kinetics but does not resolve which edited lineage drives the accelerated dissemination; lineage-resolved sampling of tissue compartments will be required to assign the phenotype to myeloid cells, CD4 T cells, or both. Finally, recent work has shown that CRISPR/Cas9-AAV6-mediated HDR editing can induce durable p53- and IL-1/NF-kB-associated inflammatory and senescence-like programs in human HSPCs, impairing long-term repopulating capacity. Although our approach relies primarily on Cas9 RNP-mediated KO rather than AAV6-based HDR, these findings highlight the need to carefully assess HSPC fitness, inflammatory activation, clonal output, and long-term engraftment during the clinical translation of edited HSPC-based therapies^68^. Genome-wide assessment of off-target editing and structural variants, which was beyond the scope of this study, will also be required before therapeutic translation.

In summary, we present a versatile HSPC-editing platform that enables efficient knockout, multiplex editing, targeted knock-in, lineage-specific functional assays, and *in vivo* HIV-1 challenge studies in humanized mice. Validation through the CCR5-KO experiment recapitulated the expected resistance to R5-tropic HIV-1, confirming the utility of this model for interrogating HIV-1 host dependency. Extending beyond proof-of-concept, the SAMHD1- KO experiment demonstrates that loss of a cellular restriction factor in the HSPC-derived human immune system accelerates HIV-1 dissemination *in vivo*. Together, these findings support the use of edited human HSPCs and humanized mice as a versatile experimental framework for dissecting HIV-1 host-factor biology and supporting the preclinical evaluation of gene- and cell-based strategies.

## Materials and Methods

### Culturing CD34^+^ HSPCs

CD34^+^ HSPCs obtained from cord blood (CB) (mixed donor) (STEMCELL Technologies), bone marrow (BM) (Lonza) and mobilized peripheral blood (MPB) (STEMCELL Technologies) were purchased and cultured in StemSpan™ SFEM II Hematopoietic Cell Culture Medium (STEMCELL Technologies) right after thawing. After nucleofection, the cells were kept in expansion medium, in which Iscove’s MDM (PAN-Biotech) was supplemented with 20% (v/v) BIT 9500 (STEMCELL Technologies), 10 µg/ml human LDL (STEMCELL Technologies), 0.1 mM 2- mercaptoethanol (Thermo Fischer Scientific), GlutaMAX (Thermo Fischer Scientific), 100 ng/ml rhFlt3-L (PeproTech), 100 ng/ml rHSPCF (PeproTech), 20 ng/ml rhIL-3 (PeproTech), 20 ng/ml rhIL-6 (PeproTech), 20 ng/ml rhG-CSF (PeproTech), 35 nM UM171 (StemCell), and 100 IU/ml Penicillin-Streptomycin (Sigma-Aldrich).

### Antibodies, cell staining and flow cytometry

For undifferentiated CD34^+^ HSPCs, megakaryocytes, and monocytes, 100 000 cells were collected. For MDMs, the cells were detached with Accutase as described above. The resultant cell suspension was washed once with PBS (Gibco) and resuspended in 50 μl FACS buffer (PBS, 1% human AB serum (Sigma-Aldrich) and 2 mM EDTA (Thermo Scientific) containing human Fc block (BD Biosciences, 1:50) and kept for 10 min on ice. Subsequently, 50 µl staining solution (FACS buffer (PBS, 2 mM EDTA, 1% FBS) and specific antibodies) was added on top of the cell suspension, which was then washed and resuspended in 100 μl FACS buffer. The following antibodies were used: anti-CD34 (pacific blue, BioLegend, cat. no. 343512, clone 581), anti- CD38 (APC, BD, cat. no. 555462, clone HIT2), anti-CD45RA (PE/Cy7, BioLegend, cat. no. 304125, clone HI100), anti -CD4 (FITC, BioLegend, cat. no. 300506, clone RPA-T4), anti-CXCR4 (APC, BD, cat. no. 555976, clone 12G5), anti-CD41 (Bv421, BioLegend, cat. no. 303730, clone HIP8), anti-CD42b (APC, BioLegend, cat. no. 303912, clone HIP1), anti-CD61 (PE/Cy7, BioLegend, cat. no. 336416, clone VI-PL2), anti-CD62P (FITC, BioLegend, cat. no. 304904, clone AK4), anti-CD46 (BV421, BD, cat. no. 743776, clone E4.3), anti PSGL-1 (AF647, BioLegend, cat. no. 328809, clone KPL-1), anti-HLA-DR (FITC, BD, cat. no. 347400, L243), anti-CCR5 (APC, BD, cat. no. 556903, 2D7). The following dyes were used: LIVE/DEAD Fixable Near-IR Dead Cell Stain kit (Thermo Fischer Scientific), LIVE/DEAD Fixable Yellow Dead Cell Stain Kit (Thermo Fischer Scientific), and CellTrace CFSE Cell Proliferation kit (Thermo Fisher Scientific) following the manufacturer’s protocol. Stained cell suspensions were analyzed with the BD FACS Lyric (BD) using FlowJo software (BD). In general, forward and side scattering of light (FSC/SSC) and LIVE/DEAD Fixable Dead Cell Stain were used to identify viable cells by flow cytometry.

### Cell proliferation assay

One day after nucleofection, WT, NTC and SAMHD1 KO cells were stained with CellTrace CFSE Cell Proliferation kit (Thermo Fisher Scientific) following manufacturer’s protocol. As a non-proliferation control, human CD4 T cells were isolated from heparinized blood retained in leukocyte reduction system chambers from healthy blood donors with approval by the Ethics Committee of the Medical Faculty of LMU München (Project No. 17-202 UE). CD4 T cells were diluted with PBS (Gibco) and CD4 T cells were isolated via the EasySep Rosette Human CD4^+^ T Cell enrichment kits (STEMCELL Technologies) according to the manufacturer’s protocol. CD4 T cells were kept in RPMI 1640 GlutaMAX (Gibco) supplemented with 10% (v/v) fetal bovine serum (FBS; Sigma-Aldrich) and Penicillin-Streptomycin (100 IU/ml; Thermo Fisher Scientific). As a proliferation control, CD4 T cells were stained with CFSE, followed by incubated with Human T-Activator CD3/CD28 magnetic beads (Gibco) and IL-2 (50 IU/ml; Biomol). CFSE fluorescence intensity was taken right after staining and every 24 h with flow cytometry.

### Colony-forming unit (CFU) assay

One day post-nucleofection, 1000 cells from NTC or SAMHD1 KO were mixed with 1 ml complete MethoCult medium (STEMCELL Technologies) and seeded in a 35 mm glass-bottom dish (ibidi), followed by incubated at 37°C for 14 days, as manufacturer’s instruction. On day 14, full-dish images were scanned with an inverted microscope (Nikon Eclipse Ti2). 3 replicate dishes were seeded for each condition, 3 full-dish images were taken for each dish. The images were exported with NIS-elements software (Nikon), and the colonies were defined as CFU-GEMM, CFU-GM, BFU-E and CFU-E manually based on the definition in Atlas of Human Hematopoietic Colonies (STEMCELL Technologies).

### KO generation in CD34^+^ HSPCs

This method is adapted from the method previously described^19^. 2 × 10^5^ CD34^+^ HSPCs (STEMCELL Technologies) were thawed and kept in culture for one day. On the next day, CD34^+^ HSPCs were washed twice with PBS (Gibco) and resuspended in 20 μl buffer P3 (Lonza; V4XP- 3032). In parallel, each synthetic sgRNA were incubated together with recombinant NLS-Cas9 (IDT; 1081059) individually for 10 min at room temperature, at a ratio of 1:2.5 (40 pmol Cas9 protein per 100 pmol gRNA) to form the 83.5 µM CRISPR/Cas9–gRNA RNP complex. 2 or 3 RNPs were used for targeting one gene in one nucleofection. For single-gene KO, 2 μl of the each RNPs were mixed with the cell suspension, and 1.25 µl of each RNPs were mixed for multi-gene KO. Cell suspension and RNPs were then transferred into a 16-well reaction cuvette of the 4D-Nucleofector System (Lonza). Cells were nucleofected using program EH-100 on the 4D-Nucleofector system. Then, 100 μl of pre-warmed IMDM Iscove’s MDM (PAN-Biotech) (without supplements) was added to each well and cells were transferred to 96-well U-bottom plates and allowed to recover for 10 min at 37 °C. Subsequently, 100 μl complete culture medium with 2 times concentrated supplements was added and the cells were kept in culture for expansion.

### Illumina MiSeq

One week after nucleofection, cells (5 × 10^4^) were collected and lysed in lysis buffer (20 μl) (1 mM CaCl_2_, 3 mM MgCl_2_, 1 mM EDTA, 1% Triton X 100 and 10 mM Tris, pH 7.5) with the addition of proteinase K (20 μg ml^−1^; Thermo Fisher Scientific). This cell lysate was incubated at 65 °C for 20 min, followed by 95 °C for 15 min and then stored at −20 °C until the PCR specific for the CRISPR/Cas9 target sites was performed. Then, 1 μl of cell lysate was used as a PCR template. For Miseq, 1 μl of lysate was used to perform PCR-I and subsequently PCR-II followed by Illumina MiSeq analysis as described previously^69^. The results of the MiSeq were analyzed with the Outknocker 2.0 webtool (http://www.outknocker.org/outknocker2.htm). For more details, please see the previous publication^19^.

### Monocyte-derived macrophage (MDM) differentiation

After nucleofection, CD34^+^ HSPCs were kept in expansion medium for 7 days, then transferred to monocyte differentiation medium for another 7 days. Monocyte differentiation medium contained IMDM Iscove’s MDM (PAN-Biotech), which supplemented with 20% FBS (Sigma-Aldrich), 20 ng/ml rHSPCF (PeproTech), 30 ng/ml rhIL-3 (PeproTech), 30 ng/ml rhFlt3-L (PeproTech), 30 ng/ml rhM-CSF (PeproTech), and Penicillin-Streptomycin (100 IU/ml; Thermo Fisher Scientific). After 7 days in monocyte differentiation medium, CD14^+^ cells were isolated with CD14 microbeads (Miltenyi Biotec) according to manufacturer’s instruction. The purity of the resultant cells was confirmed with flow cytometry and then cultured in RPMI 1640 supplemented with 10% FBS, Penicillin-Streptomycin (100 IU/ml; Thermo Fisher Scientific), and 100 ng/ml rhM-SCF, with 50000 cells each well in a 96-well plate. After 7 days, the MDMs were then subsequently used in HIV-1 infection.

### WES system western blot

Cells were lysed and the protein concentration was quantified as described above. Then, 0.6 μg of total protein was evaluated by separation and immunodetection employing the WES system (ProteinSimple) with a separation matrix of 12–230 kDa. The primary antibodies used for the WES evaluation detect SAMHD1 (proprietary mouse monoclonal antibody of the Keppler laboratory, clone H154, produced by Eurogentec; 1:200 dilution), MX2 (rabbit, polyclonal, Novus Biologicals, cat. no. NBP1-81018; 1:2000) and vinculin (mouse, hVIN-1, Sigma Aldrich, cat. no. V9264; 1:2000 dilution). The blots and quantification were generated with ProteinSimple’s Compass software (Biotechne).

### HIV-1 plasmids

The X4 tropic HIV-1 GFP proviral clone NLENG1-IRES, which is a replication competent HIV-1 NL4-3 backbone GFP reporter virus, was used and referred to as X4 HIV-1 GFP^19^. The R5 tropic NLENG1-IRES-70 was cloned from NLENG1-IRES described above by replacing env with YU-2 env. Both NLENG1-IRES and NLENG1-IRES-70 were provided by David N. Levy. For VSV-G HIV- 1 pseudovirus, VSV-G envelope expressing plasmid pMD2.G was expressed together with a construct encoding LTR-driven HIV-1 provirus NL-43 Δenv Δvpr and GFP in the nef locus.

### HIV-1 production

Sucrose cushion-purified HIV-1 stocks were produced as previously described. In brief, 293T cells were seeded at a density of 8 × 10^6^ cells in a 15-cm dish. After 24 h, cells were co-transfected with a mixture of 37.5 µg HIV-1 plasmid and 112.5 µl of L-PEI (3 µl of L-PEI for every µg of DNA; stock concentration of 1 µg/µl, Polysciences, Inc) in 2.5 ml DMEM without any additives for 30 min. After this time, 2.5 ml DNA/PEI solution was added on top of the cells. After 72 h, the supernatant was harvested and virus was purified via 25% sucrose-cushion centrifugation. For producing VSV-G HIV-1, the transfection was performed as described above, combining 37.5 μg of HIV-1 NL-43 env^−^ vpr^−^, 12.5 μg of pMD2.G and 150 µl L-PEI (1 µg/µl) for every 2.5 ml DNA/PEI solution.

### HIV-1 infection

The titer of individual virus stocks was determined on Sup T1 cells using virus-encoded GFP signals measured by flow cytometry as readout for productive infection. CD34^+^ HSPC-derived megakaryocytes and MDMs were infected with virus stocks at different MOIs as indicated for each experiment. For megakaryocytes, infections were performed by co-incubation of virus and spinoculation for 2.5 h at 650 g and 37 °C. After 3 days, cells were stained with anti-CD41 (BV421, BioLegend, cat. no. 303730, clone HIP8), anti-CD61 (PE/Cy7, BioLegend, cat. no. 336416, clone VI-PL2) and LIVE/DEAD Fixable Yellow Dead Cell Stain Kit (Thermo Fischer Scientific), followed by washed twice with PBS (Gibco) and fixed with 4% (v/v) paraformaldehyde (PFA) for 1.5 h. Cells were then washed and resuspended in FACS buffer. For MDMs, the cells were directly incubated with viruses for 3 days, without additional centrifugation. On the day of harvest, MDMs were detached with Accutase as described above. The cell suspension was washed and stained with LIVE/DEAD Fixable Near-IR Dead Cell Stain kit (Thermo Fischer Scientific), followed by 4% PFA fixation for 1.5 h. The percentage of GFP positive cells was monitored by flow cytometry. Drug were added to cells 30 min before HIV-1 challenge as control. The following drugs were used: Efavirenz (EFV, 10 µM; Sigma-Aldrich), AMD3100 (16 μg/ml; Sigma-Aldrich), and Maraviroc (MVC, 10.3 µg/ml; Sigma-Aldrich).

### IFN-α2a stimulation

After differentiating to MDMs, the cells were treated with 10 unit/ml rhIFN-α2a (Miltenyi Biotec) in RPMI 1640 medium supplemented with FBS, Penicillin-Streptomycin, and rhM-SCF as described above. One day later, the medium was replaced with fresh RPMI 1640 medium supplemented with FBS, Penicillin-Streptomycin, and rhM-SCF, and the cells were subsequently proceeding to HIV-1 infection.

### Megakaryocyte differentiation

After nucleofection, CD34^+^ HSPCs were kept in expansion medium for 3 days, then transferred to megakaryocyte differentiation medium, in which StemSpan™ SFEM II Hematopoietic Cell Culture Medium (STEMCELL Technologies) was supplemented with 50 ng/ml rhTPO-1 (PeproTech). The cells were subsequently infected with X4 HIV-1, R5 HIV-1 or VSV-G HIV-1 after 11 days in megakaryocyte differentiation medium.

### Production of knock in DNA templates

The RAB11a-GFP construct was purchased from Addgene (Plasmid #112012) and initially published by Roth et al,^18^.The DNA template was amplified by PCR using specific primers. The PCR reaction contained 5 μl 5× High-fidelity PCR buffer (Thermo Fisher Scientific), 5 μl 5× GC PCR buffer (Thermo Fisher Scientific), 1 μl dNTPs (10 mM stock; Thermo Fisher Scientific), 1.5 μl dimethylsulfoxide, 2.5 μl forward primer (10 μM stock), 2.5 μl reverse primer (10 μM stock), 1 μl Phusion (NEB), 1 μl (10 ng) plasmid and 30.5 μl H2O. The primers used for the KI template is reported in Roth et al^18^. The PCR cycle settings were 95 °C for 5 min, followed by 35 cycles at 95 °C for 30 s, 58 °C for 30 s and 72 °C for 90 s, with the final step at 72 °C for 5 min. After the PCR, a PCR clean-up was performed with the NucleoSpin Gel and PCR clean-up (Macherey-Nagel) according to the manufacturer’s protocol. Finally, the DNA concentration was determined by NanoDrop (Thermo Fisher Scientific). The GFP-RAB11A KI template is listed in Roth et al,^18^.

### Knock in of CD34^+^ HSPCs

This method was adapted from a previously published article^19^. In brief, the same nucleofection conditions as for the KO generation were used (P3 buffer and program EH-100; Lonza). In addition to the RNP, 200 ng donor DNA template was added to the P3 cell suspension. Additional information on the overall strategies to generate KIs into RAB11A loci in CD34^+^ HSPCs, including DNA templates, gRNAs and primers is included in Roth et al.^18^

### Immunoblotting

MDMs differentiated from HSPCs were incubated with Accutase solution (Sigma-Aldrich) 2 h at 37 °C and 10 min at 4 °C. The detached cells were then washed once with PBS (Gibco), followed by lysed with RIPA buffer (Cell Signaling Technology) supplemented with cOmplete™ EDTA-free Protease Inhibitor Cocktail (Roche) phosphatase inhibitors (Thermo Fisher Scientific) and kept on ice for 30 min followed by freezing at −80 °C. Cell lysates were thawed om ice and cell debris was removed by centrifugation at 10 000 g for 10 min at 4 °C. Protein concentration was quantified with BCA Protein Assay Kit (Thermo Fischer Scientific), following manufacturer’s instruction. 20 µg cell lysate were mixed with reduced LDS sample buffer (Thermo Fischer Scientific) and 50 mM dithiothreitol, followed by incubated at 90°C for 10 min. Sample lysates were separated by tris-glycine denaturing SDS-PAGE (Thermo Fischer Scientific), which were then blotted onto 0.2 mm nitrocellulose membranes (GE Healthcare). After blocking in 5% milk (Carl Roth) in TBS-T for 1 h, membranes were incubated with the following primary antibodies overnight at 4 °C: anti-RAB11A (rabbit, polyclonal, Thermo Fischer Scientific, cat. no. 71-5300) 1:1 000 dilution, anti-GFP (rabbit polyclonal, Chromoteck, cat. no. PABG1-20) 1:1 000 dilution, and anti-vinculin (mouse, hVIN-1, Sigma-Aldrich, cat. no. V9264) 1:2 000 dilution. HRP-conjugated goat anti-mouse IgG (rat adsorbed, polyclonal, Bio-Rad, cat. no. STAR77), and goat anti-Rabbit IgG (H + L, polyclonal, Dianova, cat. no. AFK-600) were used in a dilution of 1:10 000. ECL (ThermoFisher Scientific) was used as substrate and the chemiluminescent signals were detected on a Fusion Fx (Vilber).

### Confocal microscopy

CB CD34^+^ HSPCs were differentiated into megakaryocytes for 14 days, and the cells were collected and stained with anti-CD42b (APC, BioLegend, cat. no. 303912, clone HIP1) and anti- CD62P (FITC, BioLegend, cat. no. 304904, clone AK4), followed by fixing in 4% (v/v) PFA. Fixed cells were mounted with ProLong Diamond Antifade Mountant with DAPI (Thermo Fischer Scientific). Images were taken with a spinning disc confocal microscope (CSU-W1, Nikon).

### Humanized mice studies

All animal experiments were reviewed and approved by the Cantonal Veterinary Office of Zurich, Switzerland (cantonal approval number ZH081/2021 and national approval number 33654) and performed in accordance with local guidelines and Swiss animal protection law. The use of Cord blood samples was covered by KEK-StV Nr. 40/14. Humanized (hu) -mice were generated as previously described by Audige et al.^70^. In brief, immunodeficient NOD.Cg-Prkdcscid Il2rgtm1Wjl/SzJ (NSG) mice were obtained from Charles River Laboratories and bred and maintained at the Laboratory Animal Service Center of the University of Zurich. Newborn mice were irradiated with 1 Gy for 37 sec 1-3 days after birth and subsequently transplanted with 1.5 ± 0.5x10^5^ CD34^+^ cells via intrahepatic injection. We checked the degree of human cell engraftment at approximately 10-12 weeks of age by staining peripheral blood for the pan-human marker CD45. Hu mice with human engraftment level of CD45^+^ cells > 5% were used for further experiments. For HIV infected mice, an intraperitoneal injection at a media tissue culture infectious dose of 2 x 10^5^ with HIV-1 YU2 was done as previously reported^71^.

### Mouse tissues preparation for flow cytometry and ex vivo analysis

For *ex vivo* analysis mice blood and spleen mononuclear cells were isolated by density gradient centrifugation using Lymphoprep (StemCell Technologies, Vancouver, Canada, catalog no. 18060), after smashing the organ on 70-μm cell strainers. Red blood cells were removed using ACK lysing buffer (Lonza, Basel, Switzerland, cat.no. 10-548E). Cells were washed twice in magnetic-activated cell sorter buffer phosphate-buffered saline (PBS) with 2 mM EDTA and 2% FBS] and stained with anti-human CD45 PerCP, CD3 BV786, CD4 PE-Cy7, CD8 BV421, CD19 PE, CD14 PE-Dazzle, CD56 BV605. Samples were fixed with 1% PFA for 40 min at 4 °C before analysis at the BD LSR Fortessa Cell Analyzer (BD Biosciences, Franklin Lakes, NJ, USA).

### Mouse tissue preparation and staining for immunofluorescence

Organs were harvested for histological analyses. Tissues were fixed in 10% formalin overnight and then stored in 1%formalin-PBS.

### Mouse sera HIV-1 viral load determination

Mouse sera were diluted 1:80 in 0.9% NaCl, and viral load was determined using either the Abbott RealTime HIV-1 Assay (Abbott Molecular, art. no. 02G31-010) on the semi-automated Abbott m2000sp/m2000rt platform or the cobas® HIV-1 assay (Roche Diagnostics, art. no. 09 040 803 190) on the fully automated Roche cobas® 6800 system, according to the manufacturer’s instructions, with the exception of the sample matrix. Testing was performed in the ISO 15189-accredited routine diagnostics laboratory of the Max von Pettenkofer Institute, Virology department, LMU Munich. Low-positive, high-positive, and negative controls were included in each run.

## Supporting information

Supplementary informations

## Acknowledgments

This work was supported by the Deutsche Forschungsgemeinschaft: grant KE742/4-2 (to O.T.K.), a grant as part of SPP-1923 (to O.T.K.), The Deutsche Zentrum fuer Infektionsforschung (DZIF), project TTU 04.820 (to O.T.K.), LMUexcellent: M.A., and the Friedrich-Baur- Foundation: Young scientist grant to M.A. and H.R. C., MFAG AIRC-28809 (M.A.), and FIS2 (MUR) FIS-2023-00896, Postdoctoral Research Abroad Program (Ministry of Science and Technology, Taiwan) (to H.R. C.) and UZH Global Strategy and Partnerships Funding Scheme, University of Zurich, Switzerland (grant do R.F.S.).

## Notes

### Competing Interest Statement

The authors have declared no competing interest.

