## Supplementary informations for "Optimizing CRISPR/Cas9 genome editing in primary human hematopoietic cells to advance studies into HIV biology"

**Supplemental Figure 1**

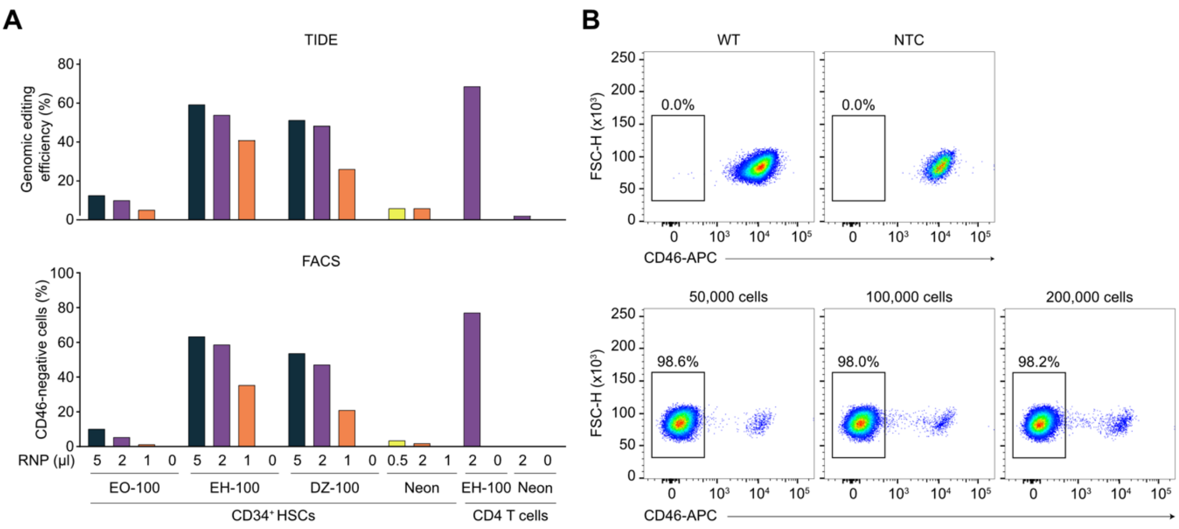


**Supplemental Figure 1. Optimizing the nucleofection for KO CD46 in CD34$^{+}$ HSCs.** (A) CD46 KO was carried out with different amounts of single-gRNA RNP complex and different programs on 4D-nucleofector or Neon Transfection System. Primary resting CD4 T cells were used as a control. For each condition, 50,000 CD34$^{+}$ HSCs or 2,000,000 CD4 T cells were used. 5 days after electroporation, cells lysate was collected for checking KO efficiency with TIDE analysis (upper panel). 7 days after electroporation, CD46 protein level was assessed by flow cytometry (lower panel). (B) CD46 KO was carried out on CD34$^{+}$ HSCs with 5 µl RNPs containing 2 sgRNAs targeting CD46 by using program EH-100 on 4D-nucleofector. 50,000, 100,000, or 200,000 CD34$^{+}$ HSCs were used per nucleofection. 7 days after nucleofection, CD46 level was assessed by flow cytometry.

**Supplemental Figure 2**

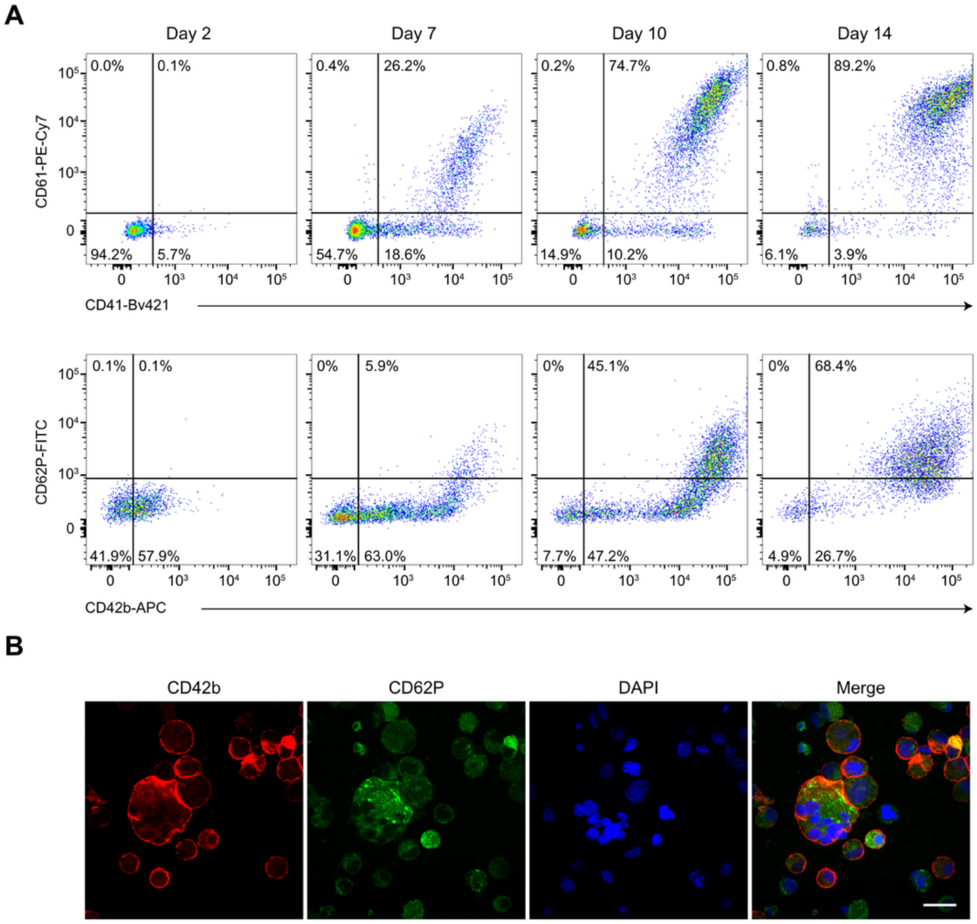


**Supplemental Figure 2. CB CD34⁺ HSCs were differentiated into megakaryocytes after 14 days in culture.** (A) Wildtype CD34⁺ HSCs were cultured in megakaryocyte differentiation medium for 14 days and the differentiation was evaluated by the expression of early megakaryocyte markers CD41 and CD61, and mature megakaryocyte markers CD42b and CD62P with flow cytometry on day 2, 7, 10 and 14. The plots of one donor are shown. (B) After 14 days, differentiated megakaryocytes were stained with CD42b and CD62P, followed by fixation with 4% paraformaldehyde and mounted with anti-fade mounting medium containing DAPI. One representative image is shown. Scale bar = 20 µm.

**Supplemental Figure 3**

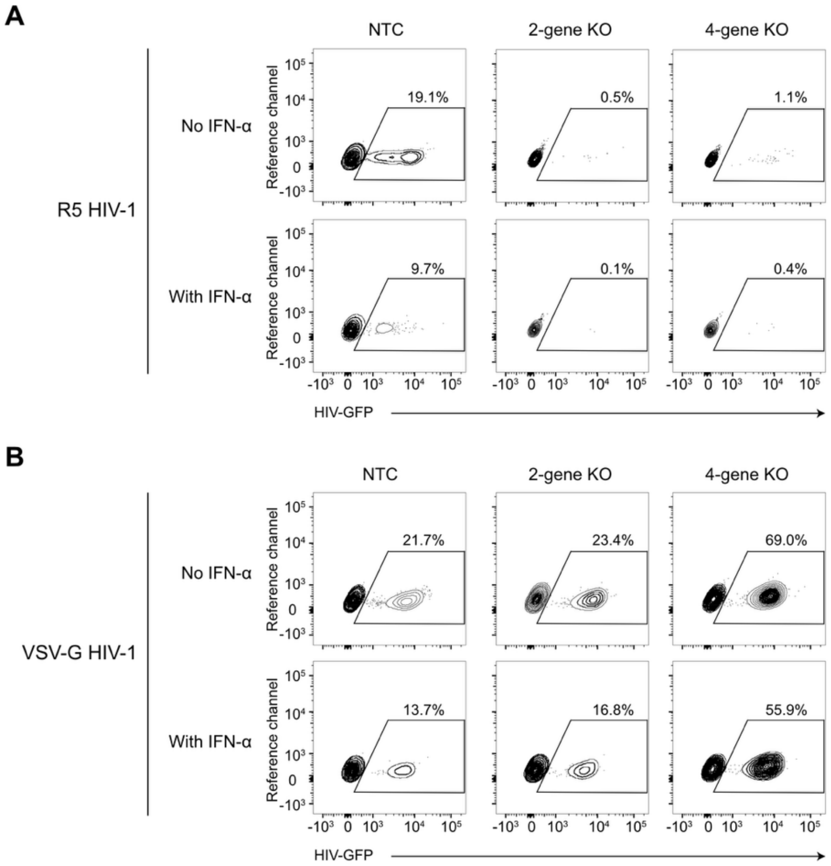


**Supplemental Figure 3.** (Density plots of Figure 5E and F). CB CD34$^{+}$ HSCs were nucleofected with RNPs targeting NTC or CD4, and CCR5 (2-gene KO), or CD4, CCR5, MX2, and SAMHD1 (4-gene KO), followed by MDM differentiation and HIV-1 infection as described in Figure 5E. The flow cytometry density plot of one representative donor is shown. (A) The density plots of R5 HIV-1 infection. (B) The density plot of VSV-G HIV-1 infection.
